# A detailed dissection of the expression, localization, structure, and diagnostic potential of cyst wall proteins of the eye pathogen *Acanthamoeba*

**DOI:** 10.1101/2024.02.02.578540

**Authors:** Bharath Kanakapura Sundararaj, Manish Goyal, John Samuelson

## Abstract

The cyst wall of the eye pathogen *Acanthamoeba castellanii* contains cellulose and chitin and has ectocyst and endocyst layers connected by conical ostioles. Previously, we used mass spectrometry of purified walls to identify an abundant laccase and three families of lectins (Jonah, Luke, and Leo). Here we show that frameshifts in the protein prediction in AmoebaDB, which incorrectly add 12 transmembrane helices, cause Jonah to mislocalize to a ring around ostioles rather than to the ectocyst layer. RT-PCR, double labels with GFP and RFP or mCherry, and promoter swaps show that ectocyst localization does not just correlate with but is caused by earlier expression, while localization in the endocyst layer and ostioles is caused by later expression. A chitin-binding domain from an *Entamoeba* chitinase shows chitin forms thick fibrils in the ectocyst layer and a honeycomb in the endocyst layer. AlphaFold shows Ac wall proteins originate from bacteria by horizontal gene transfer (β-helical folds of Jonah and three cupredoxin-like domains of the laccase), share common ancestry with wall proteins of slime molds (β-jelly-roll folds of Luke), or are unique to *Acanthamoeba* (four disulfide knots of Leo). Ala mutations show linear arrays of aromatic amino acids in β-jelly-roll folds of Luke and disulfide knots of Leo are necessary for binding cellulose and proper localization of proteins in the cyst wall. Finally, rabbit antibodies to recombinant Jonah, Luke, Leo, and laccase efficiently detect calcoflour white-labeled cysts of 10 of 11 *Acanthamoeba* isolates tested, suggesting all four proteins are excellent diagnostic targets.

**IMPORTANCE:** *Acanthamoebae* are free-living amoeba in the soil and water that cause *Acanthamoeba* keratitis in under-resourced countries, where water for washing hands may be scarce. *Acanthamoeba* is an emerging pathogen in the United States, because of its association with contact lens use. Here we show early expression during encystation causes a Jonah lectin and a laccase to localize to the outer layer of the cyst wall, while later expression cause Luke and Leo lectins to localize to the inner layer and the conical ostioles that connect the layers. We used structural predictions to identify the aromatic amino acids of Luke and Leo necessary for binding cellulose in the wall and to identify domains of Jonah and laccase useful for making recombinant proteins to immunize rabbits. Rabbit antibodies to Jonah, Luke, Leo, and laccase all efficiently detected cysts of ten *Acanthamoeba* isolates, including five T4 genotypes that cause most keratitis cases.

## Introduction

*Acanthamoeba* keratitis (AK), which leads to scarring and blindness, if not successfully treated, is caused by free-living amoebae, named for acanthopods (spikes) on the surface of trophozoites (1–3). In immunocompromised patients, *Acanthamoebae* may cause encephalitis, which is frequently fatal (4). AK is associated with corneal trauma in the Middle East, South Asia, and Africa, where water for handwashing is often scarce (5–9). AK is associated with contact lens use in the US, Europe, and Australia and so *Acanthamoeba* is on the NIAID list of Emerging Infectious Diseases/Pathogens (3, 10–12). Regardless of the place or the cause of infection, 18S RNA sequences show that the T4 genotype most often causes AK, while other genotypes less frequently or never cause AK (13–16). The whole genome of the Neff strain of *Acanthamoeba castellanii* (Ac), a T4 genotype that is broadly studied, was sequenced, and proteins were predicted and deposited in AmoebaDB (17, 18). Recently, the Neff genome has been re-sequenced with long reads to produce 33 chromosome-size assemblies, while protein prediction has been improved by transcriptomes of trophozoites and organisms encysting for eight hours (19, 20).

Cyst walls, which form when trophozoites are starved of nutrients, protect free-living *Acanthamoebae* from osmotic shock in freshwater or drying in the air (21, 22). The cyst wall also makes *Acanthamoebae* resistant to disinfectants used to clean surfaces, alcohol-based hand sanitizers, sterilizing agents in contact lens solutions, and/or antibiotics applied to the eye (23–27). While trophozoites cause damage to corneal epithelial cells, AK is most often diagnosed by microscopic identification of cysts in corneal scrapings (1, 2, 28). *Acanthamoeba* cysts are detected using calcofluor white (CFW) or wheat germ agglutinin (WGA), which bind to cellulose and chitin, respectively, in their walls (29, 30). Because CFW and WGA also react with walls of plants, fungi, and other protists, they are suboptimal reagents that may lead to incorrect diagnosis of AK (31–33). The translational goal of the present study, therefore, is to identify abundant cyst wall proteins, which might be targets for diagnostic antibodies. Antibodies to the *Acanthamoeba* wall might also be useful to detect cysts in concentrated water samples, monitor contamination in contact lens solutions, test reagents for cleaning reusable contact lenses, and/or test drugs that inhibit encystation.

Nearly 50 years ago, cellulose (β-1,4-linked glucose) was identified in *Acanthamoeba* cyst walls, which has an outer ectocyst layer and an inner endocyst layer connected by conical ostioles (34, 35). Because *Acanthamoeba* has a chitin synthase and WGA binds to cyst walls, it is also likely that chitin (β-1,4-linked GlcNAc) is also present (36). No cyst wall proteins were known until we purified cyst walls from the Neff strain of *Acanthamoeba castellanii* (Ac), which is a T4 genotype, and identified three families of cellulose-binding lectins that we named Jonah, Luke, and Leo (29). When a representative protein from each family was tagged with green fluorescent protein (GFP) and expressed under its own promoter in transfected Ac, Jonah localized to the ectocyst layer that is made early, while Luke and Leo localized to the endocyst layer and ostioles that are made later (37, 38). Anti-GFP antibodies showed Jonah-GFP is more accessible than Luke-GFP and Leo-GFP, suggesting Jonah and other ectocyst proteins might be better targets for anti-cyst antibodies to diagnosis AK by fluorescence microscopy of eye scrapings (29).

Here we performed basic science experiments to understand better the localization, origin, and structure of Jonah, Luke, and Leo lectins, as well as an abundant laccase, as well as translational science experiments to test their value as targets for antibodies to diagnose cysts in patients with AK. First, we identified three additional Ac proteins in the ectocyst layer and used RT-PCR, double labels with GFP and RFP or mCherry, promoter swaps, and a GFP-tagged probe for chitin to begin to understand why proteins are located in the ectocyst layer versus the endocyst layer and ostioles. Second, we used AlphaFold and Foldseek to better understand the structure and origin of wall proteins (39–41), and we made alanine mutations to show linear arrays aromatic amino acids of Luke and Leo bind cellulose (42–47). Third, we tested rabbit antibodies (rAbs) to recombinant Ac cyst wall proteins against cysts of 10 other *Acanthamoeba* strains and species, including five T4 genotypes that cause most cases of AK (13–16).

## RESULTS

### Localization of Jonah-3, Leo-S, and laccase-1 to the ectocyst layer of mature walls of Ac

The goal here was to identify additional abundant Ac cyst wall proteins that localize to the ectocyst layer, which is likely more accessible to diagnostic antibodies than the endocyst layer. Jonah family lectins are highly abundant in cyst walls and have an N-terminal signal peptide followed by one or three choice-of-anchor A (CAA) domains, which form β-helical folds (BHFs) (Fig. 1A). Jonah-1 (ACA1_164810), which has a long, unstructured Thr-rich domain in addition to a single BHF, is expressed early during encystation and localizes to the ectocyst layer when tagged with GFP and expressed under its own promoter in transformed Ac of the Neff strain (ATCC 30010) (Fig. 1A, 1B, and 1E) (Supporting Information S1 and S2) (29, 37–41). The ectocyst layer is labeled with WGA, which binds to chitin, while the endocyst layer and ostioles are labeled with CFW, which binds to cellulose and/or chitin. Here we tested the localization of Jonah-3 (ACA1_157320), which contains three BHFs and was very abundant in purified cyst walls (17, 18, 29). While other cyst wall proteins contain a signal peptide but no TMHs, Jonah-3-AmoebaDB contains 12 transmembrane helices (TMHs) (Fig. 1A) (48, 49). GFP-tagged Jonah-3-AmoebaDB, which was expressed under its own promoter, localizes to punctate rings around each ostiole (Fig. 1B and 1C). To test whether this ring-like localization of Jonah-3-AmoebaDB might be an artifact of incorrect protein prediction, we used an unfinished transcriptome of encysting Ac to make Jonah-3(c) (corrected), which contains three BHFs separated by unstructured, Ser-rich spacers (Fig. 1A and 1F and Supporting Information S1). An alignment of Jonah-3(c) and Jonah-3-AmoebaDB shows 12 TMHs in the latter derive from a series of frameshifts rather than deletions or insertions. Further, Jonah-3(c)-GFP localizes to the ectocyst layer with an identical appearance to that of Jonah-1-GFP (Fig. 1D).

**FIG 1.**
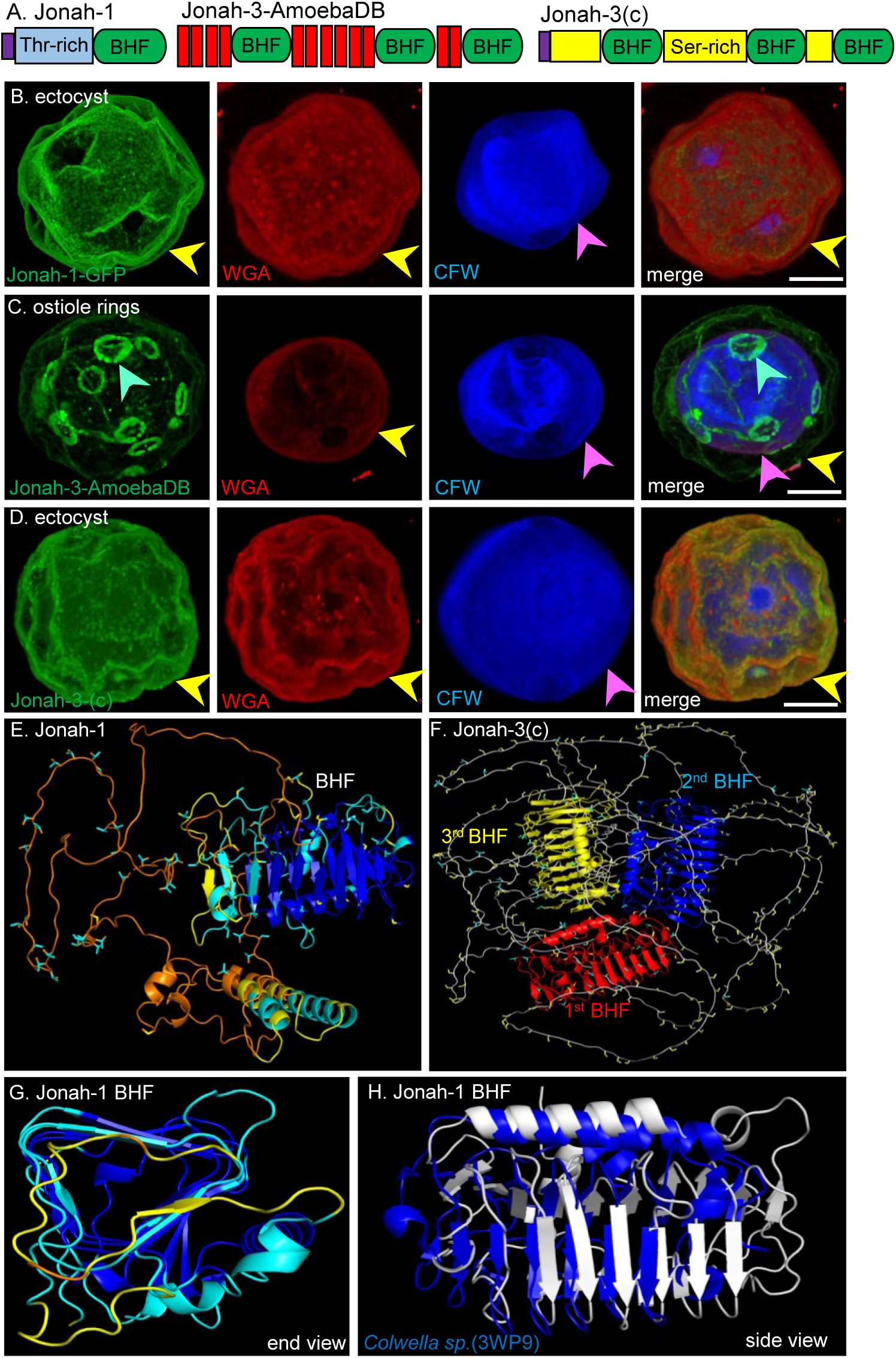
Localization and structure of Jonah lectins. (A) Drawings show Jonah-1 contains a signal peptide (purple), followed by a Thr-rich spacer (light blue), and a single BHF (green). Jonah-3-AmoebaDB has twelve TMHs (red) and three BHFs, while Jonah-3(c) has three BHFs, three Ser-rich domains (yellow), and no TMHs. (B) Confocal microscopy shows Jonah-1 localizes to the ectocyst layer (yellow arrows, stained with WGA). The endocyst layer is marked with CFW (pink arrow), while ostioles appear as dimples (purple arrows). (C) Jonah-3-AmoebaDB with 12 TMHs forms rings around ostioles (green arrows) when expressed under its own promoter with a GFP tag. (D) Jonah-3(c) localizes to the ectocyst layer, like Jonah-1. Confocal micrographs here and in Fig 2 to 7 were shot with a 60x objective, and 3D reconstructions were made from 0.1 µm optical sections. (E) AlphaFold structure with confidence colored shows the Jonah-1 lectin is composed of a single three-sided, β-helical fold (BHF), as well as two α-helices (function unknown), and a 170-aa, unstructured domain rich in Thr (light blue). (F) Jonah-3-(c) has three BHFs (red, blue, and yellow) and 132-aa, 267-aa and 308-aa long, unstructured domains rich in Ser (yellow). (G) End-view with confidence colored shows the Jonah-1 BHF is three-sided (BHFs may also be two-sided). (H) Foldseek analysis shows the Jonah-1 BHF (white) closely matches the BHF of *Colwella sp.* (blue), the structure of which has been solved (PDB 3WP9). The E-value is 5.10e-4, and the RMSD is 9.21, despite a sequence identity of just 12% over a 245-aa overlap. Scale bars for B to D are each 5 µm.

Luke-2 (ACA1_377670) and Luke-3 (ACA1_245650), which have two or three β-jelly-roll folds (BJRFs), respectively, that are separated by unstructured, Ser-rich spacer(s), are expressed later during encystation and localize to the endocyst layer and ostioles when tagged with GFP and expressed under their own promoters (Fig. 2A, 2B, 2D, and 2E). Leo-A (ACA1_074730), which contains two adjacent sets of four disulfide knots (4DKs), is also expressed later in encystation and localizes to the endocyst layer and ostioles when tagged with GFP (Fig. 3A, 3B, and 3G). Here we tested the localization of Leo-S lectin (ACA1_188350), which contains two sets of 4DKs separated by a long, unstructured, Thr-rich spacer (Fig. 3A and 3I). We used for a second time the unfinished Ac transcriptome to make Leo-S(c) that corrects an eight amino acid deletion in the N-terminal 4DK, which was identified by an alignment with other Leo-S lectins (Supporting Information S1). Unlike Leo-A-GFP, which localizes to the endocyst layer and forms a flat ring around the ostioles, Leo-S(c)-GFP localizes to the ectocyst layer of mature cyst walls in a pattern like those of Jonah-1 and Jonah-3(c) (Fig. 3F). Time course studies of Leo-S(c)-GFP during the encystation showed that after 12 hours encystation, Leo-S(c)-GFP is present in a dense set of secretory vesicles (SV in Fig. 3D) that fill the cytosol of cells, which are rounded but lack a wall (as shown by a failure to label with WGA or CFW). However, after 24 hours encystation, Leo-S(c)-GFP forms a patchy distribution on a single-layered wall, which is labeled with both WGA and CFW (Fig. 3E). After 48 hours, Leo-S(c)-GFP has a homogeneous distribution in the ectocyst layer (data not shown), which is the same as Leo-S(c)-GFP in mature cysts at 72 hours (Fig. 3F).

**FIG 2.**
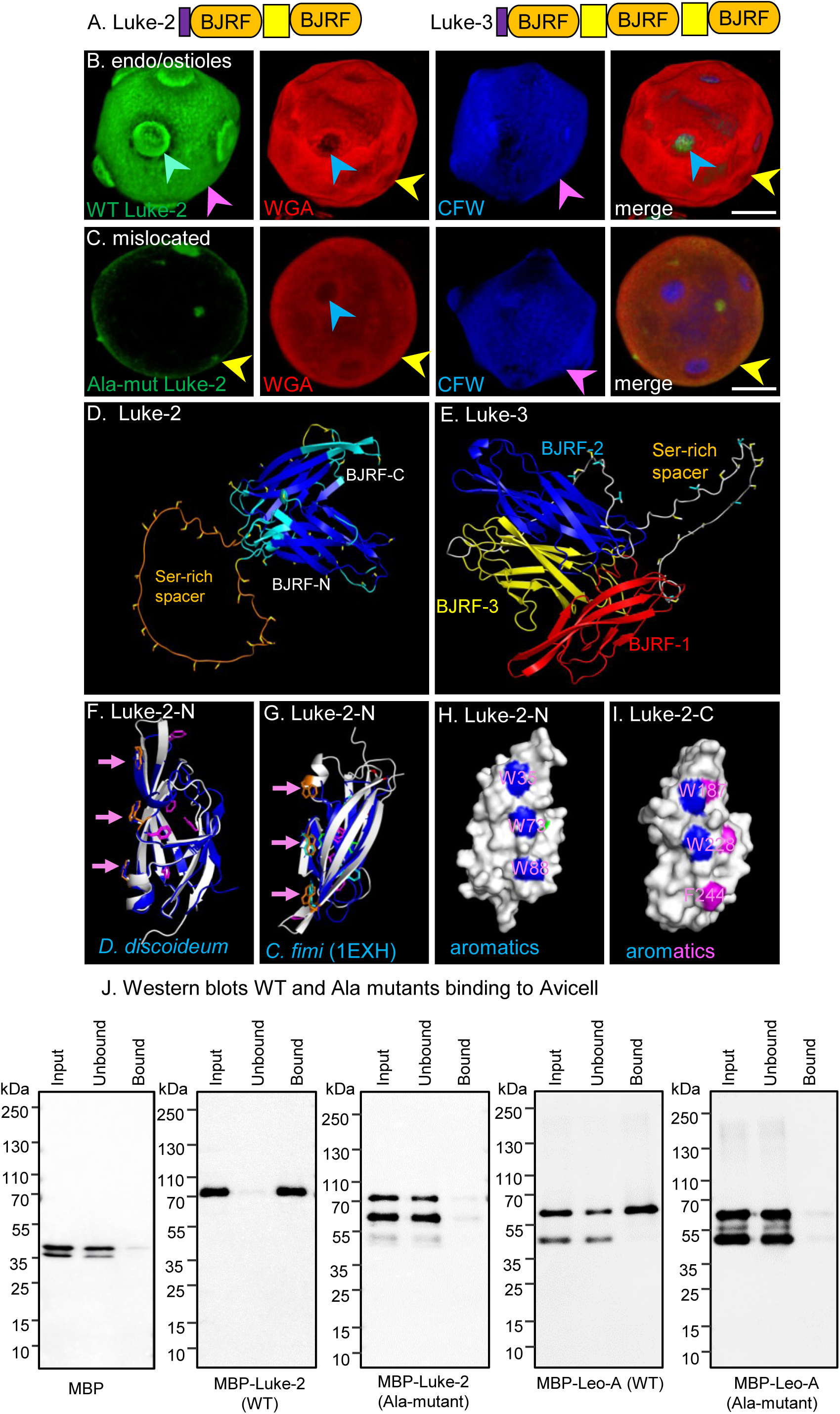
Structure and localization and structure of Luke-2. (A) Drawings show Luke-2 and Luke-3 contain a signal peptide (purple) and two or three β-jelly roll folds (BJRFs) (orange), respectively, which are separated by Ser-rich spacers (yellow). (B) Confocal microscopy shows WT Luke-2-GFP localizes to the endocyst layer (pink arrows) and forms a flat ring around ostioles (blue-green arrows) (see also Fig. 5D). (C) When five Tyr and a Phe are mutated to Ala (see H and I below), Ala-mut Luke-2-GFP no longer localizes to endocyst layer. (D) AlphaFold shows Luke-2 contains two BJRFs, which are separated by a 47-aa unstructured, spacer rich in Ser (yellow). (E) Luke-3 has three BHFs separated by 42-aa and 33-aa long, unstructured spacers that are also Ser-rich. (F and G). Foldseek shows the N-half BJRF of Luke-2 shares a similar structure with a BJRF of *Dictyostelium* wall protein and with CBM2 of a *Cellulomonas fimi* endoglucanase (blue), the structure of which has been solved (PDB 1EXH). The E-value for the BJRFs of Luke-2 and *D. discoideum* wall protein is 4.99e-7, and the RMSD is 1.79, strongly suggesting common ancestry. The E-value for the N-terminal BJRF of Luke-2 and the CBM2 of *C. fimi* is 9.72e-5, and the RMSD is 5.01, despite a 16% identity over a 105-aa overlap. The N-terminal BJRF of Luke-2 has three linearly arranged aromatics (orange), which match those of *D. discoideum* and *C. fimi* (pink arrows). H and I. Surface views of Luke-2-N and Luke-2-C highlight linear arrays of Trp (blue) and Phe (pink), which were mutated to make Ala-mut of Luke-2-GFP that no longer localizes to the endocyst layer and ostioles (see B above). J. Western blots show that MBP alone fails to bind to Avicel cellulose, while MBP-Luke-2 and MBP-Leo-A fusion-proteins (described in Fig. 3) each bind well to Avicel cellulose. In contrast, cellulose binding is lost when six aromatics are mutated to Ala in both Luke-2 and Leo-A. These experiments support the idea that Ala-mutants of Leo-A and Luke-2 mislocalize in cyst walls, because of the loss of cellulose binding. Scale bars for B and C are each 5 µm.

**FIG 3.**
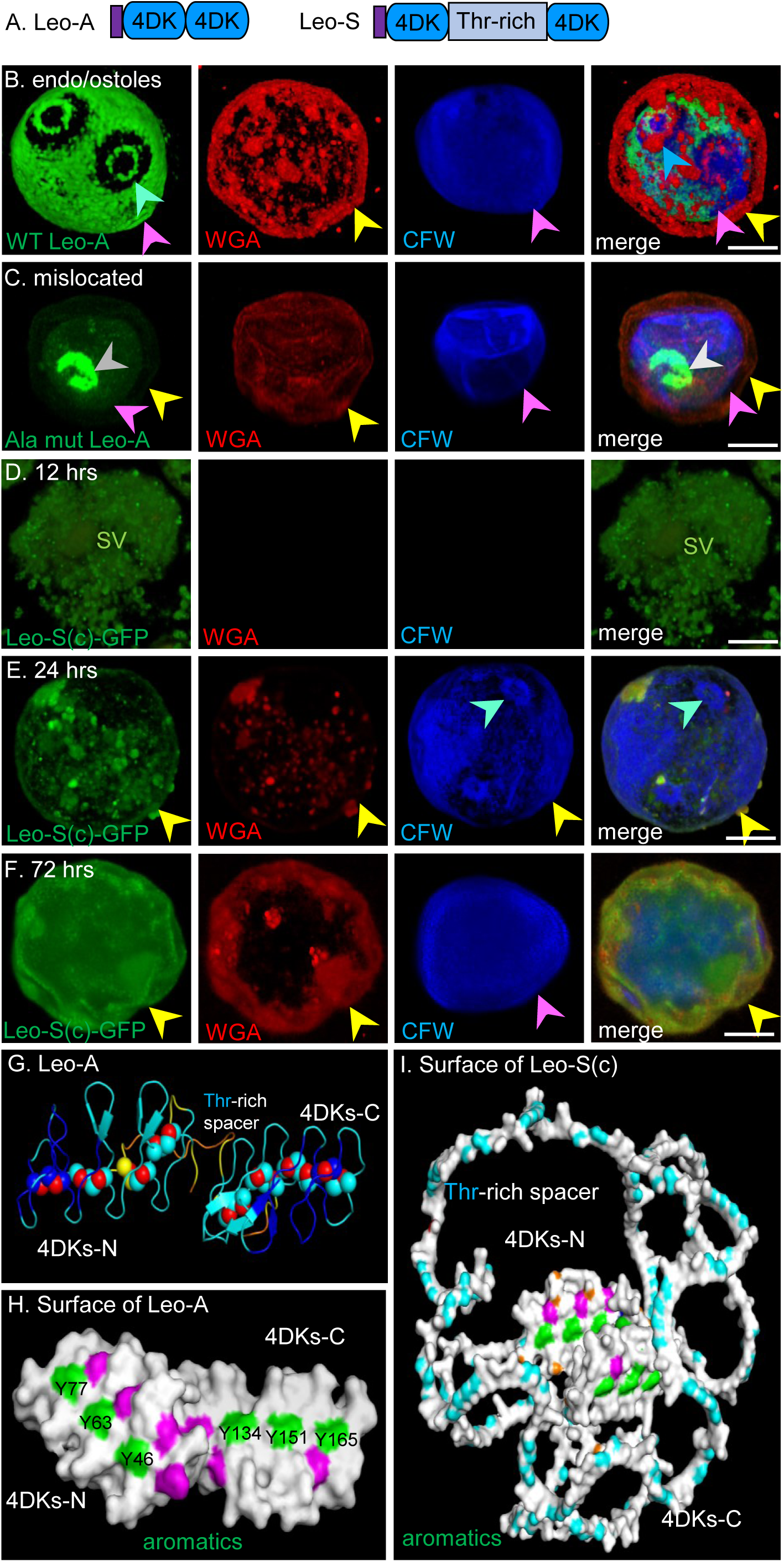
Structure and localization of Leo lectins. (A) Drawings show Leo-A contains a signal peptide (purple) and two adjacent sets of four disulfide knots (4DKs) (blue), while Leo-S contains a Thr-rich spacer (grey-blue) between sets of 4DKs. (B) Confocal microscopy shows WT Leo-A localizes to the endocyst layer (pink arrows) and forms a flat ring around ostioles (blue-green arrows) (see also Fig 5D). (C). When six Tyr are mutated to Ala (see H below), Ala-mut Leo-A (grey arrow) no longer localizes to endocyst layer. (D) Early in encystation (12 hours), secretory vesicles (SV) containing Leo-S(c) fill the cytosol of Ac, which lack a cyst wall as shown by failure to label with WGA or CFW. (E) Later in encystation (24 hours), Leo-S(c) has a somewhat patchy distribution in the single-layered wall, which now labels with WGA and CFW. (F) In mature cyst walls (72 hours), Leo-S(c) has a homogeneous distribution in the ectocyst layer (yellow arrows). Note that CFW predominantly labels the endocyst layer of mature cysts. (G) A ribbon diagram of Leo-A highlights 4DKs (red spheres), which connect short, parallel loops. (H) The surface of Leo-A shows three Tyr residues (green) linearly arrayed in each 4DK, which were mutated to Ala to test the effects on localization (see C above) and cellulose-binding (see Fig 2J). (I) The surface of Leo-S-corrected shows a 252-aa long unstructured, spacer rich in Thr (blue) between 4DKs, each of which contains linearly arrayed Tyr residues. Foldseek had no hit with the structure of Leo-A. Scale bars for B to F are each 5 µm.

The last protein tested was laccase-1 (ACA1_068450), which is a multicopper oxidase with three cupredoxin-like domains (CuRO-1, CuRO-2 and CuRO-3) that is abundant in purified cyst walls of Ac (Fig. 4A and 4E) (29, 50, 51). Laccase-2(c) (ACA1_006180), the sequence of which we corrected using the unfinished transcriptome, is much less abundant and so was not localized. GFP-tagged laccase-1 under its own promoter localizes to the ectocyst layer of mature Ac walls with an appearance like those of GFP-tagged Jonah-1, Jonah-3(c), and Leo-S(c) (Fig. 4D). Like Leo-S(c)-GFP, laccase-1-GFP fills secretory vesicles of Ac encysting for 18 hours, when there is no wall labeled with WGA or CFW (Fig. 4B). N.B. that the precise timing of secretory vesicles and wall formation may vary by up to 6 hours, depending on the exact condition of trophozoites prior to placement in encystation medium. Unlike Leo-S(c)-GFP, after 24 hours encystation laccase-1-GFP is homogeneously distributed in the single layered wall of Ac, which becomes the ectocyst layer of mature cysts at 72 hours (Fig. 4C and 4D).

**FIG 4.**
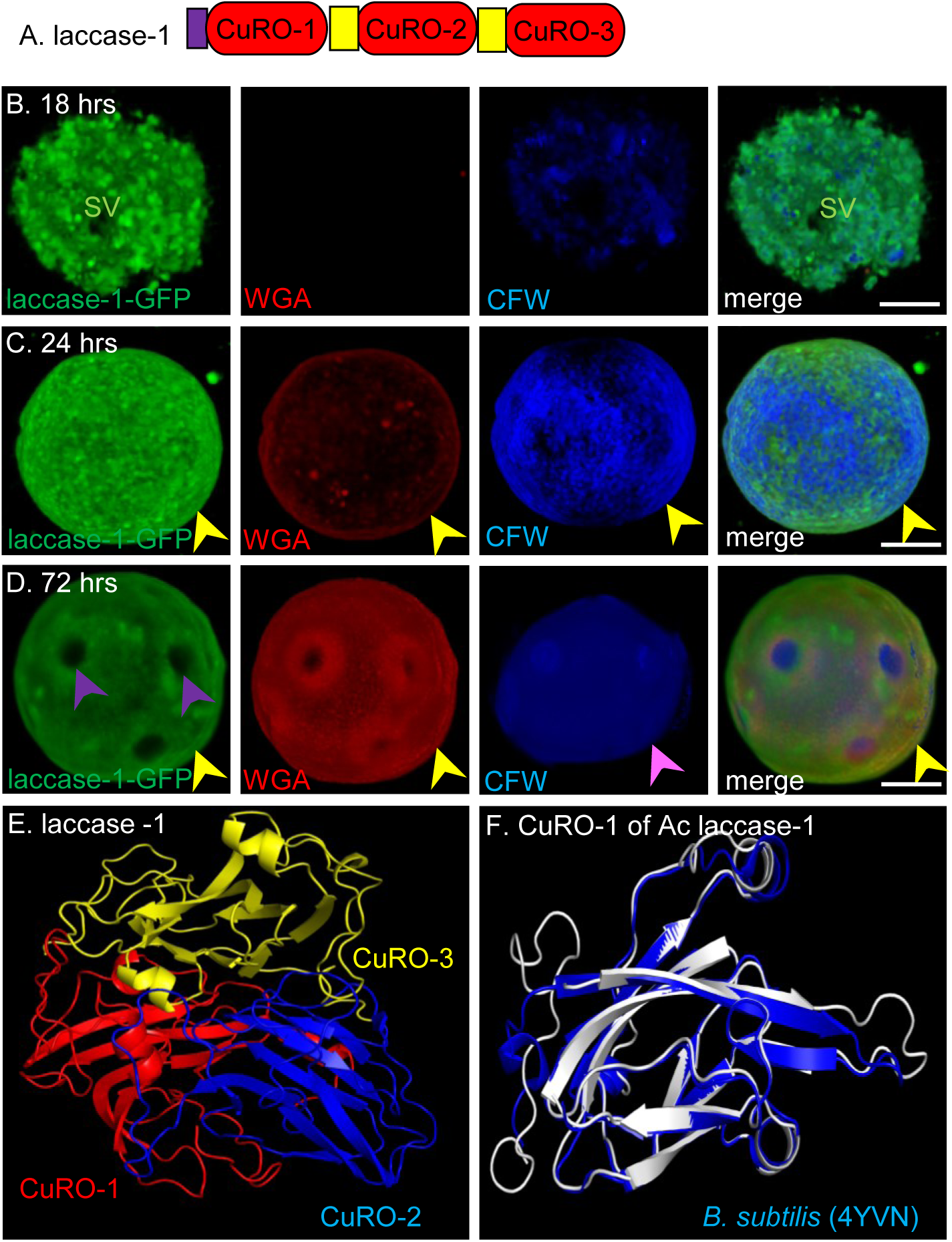
Localization and structure of an abundant laccase. (A) The Ac Laccase-1 has signal peptide (purple) and three copper oxidase domains (red), separated by short, unstructured spacers lacking either Ser- or Thr (white). (B) Confocal microscopy shows that early in encystation (18 hours) secretory vesicles (SV) containing laccase-GFP fill the cytosol of Ac, which lack a cyst wall as shown by failure to label with WGA or CFW. (C) Later in encystation (24 hours), laccase-GFP has a homogeneous distribution in the single-layered wall, which now labels with lightly with WGA and somewhat more with CFW. (D) In mature cyst walls (72 hours), laccase-GFP has the same homogeneous distribution in the ectocyst layer (yellow arrows) with ostioles as dimples (purple arrows). (E) AlphaFold structure shows CuRO-1 (blue), CuRO-2 (red), and CuRO-3 (yellow) domains of the laccase, all of which are predicted with confidence (not shown). (G) Typical for an enzyme, the E-value is 1.32e-59 and the RMSD = 2.75 with a sequence identity of 39% over a 505-aa overlap. Scale bars for B to D are each 5 µm.

In summary, we localized three new proteins to the ectocyst layer of the Ac cyst wall, one of which was correctly predicted by AmoebaDB (laccase-1) and two of which needed to be corrected using an unfinished transcriptome (Jonah-3(c) and Leo-S(c)). We also showed that Leo-S(c) and the laccase-1 are each made early during encystation in a dense set of secretory vesicles, which are released onto the single-layered wall in a fashion like that of Jonah-1 (29).

### RT-PCR and promoter swaps show timing of expression does not just correlate with but causes protein localization in the ectocyst layer (earlier) or the endocyst layer and ostioles (later)

Four experiments here address limitations in our present understanding of how proteins are targeted to the two layers of the Ac cyst wall and ostioles. First, we use qRT-PCR to track mRNAs for Jonah-1, Luke-2, Leo-A, and laccase-1 to determine the extent to which the results correlate with confocal micrographs of GFP-tagged proteins each expressed under its own promoter (∼500-bp upstream of the start ATG) (Fig. 4B to 4D) (29). Second, to visualize the development of the two layers of the wall in the same organism, we encysted Ac expressing Luke-2-GFP and either Jonah-1-RFP or Jonah-1-mCherry, each under its own promoter on a single plasmid, and examined them with confocal microscopy plus CFW. Third, we use promoter swaps between pairs of proteins (Jonah-1 and Luke-2 or laccase-1 and Leo-A) to test the hypothesis that early expression does not just correlate with but causes Ac cyst wall proteins to localize to the ectocyst layer, while later expression causes cyst wall proteins to localize to the endocyst layer and ostioles (29). We also tested here whether proteins expressed under the same promoter have the similar or different localizations in the Ac cyst wall, which would suggest similar or different binding specificity, respectively, for cellulose and/or chitin, the two glycopolymers that form fibers in the wall (35, 36). Fourth, we use earlier and later promoters (Jonah-1 and Luke-2, respectively) to express a heterologous probe for chitin fibrils, which is composed of a chitin-binding domain of an *Entamoeba* chitinase called CBM55 fused to GFP (52).

Across nine time points, we isolated RNA from nontransfected trophozoites and Neff strain encysting for 96 hours and performed qRT-PCR with cDNA (Fig 5A). qRT-PCR products were normalized by calreticulin, which is part of the *N*-glycan-dependent quality control of protein folding in the ER and is highly expressed in trophozoites and encysting Ac (53). Transcription of Jonah-1 begins at 6 hrs encystation and peaks at 18 hrs, while transcription of laccase-1 begins at 12 hrs and peaks at 24 hrs. In contrast, transcription of Luke-2 begins at 24 hrs and peaks at 48 hrs, while transcription of Leo-A is only detected at 72 hrs. The qRT-PCR results agree with those of the unfinished transcriptome, which also suggests the low level of Leo-A is likely caused by a suboptimal choice of primers for RT-PCR.

**FIG 5.**
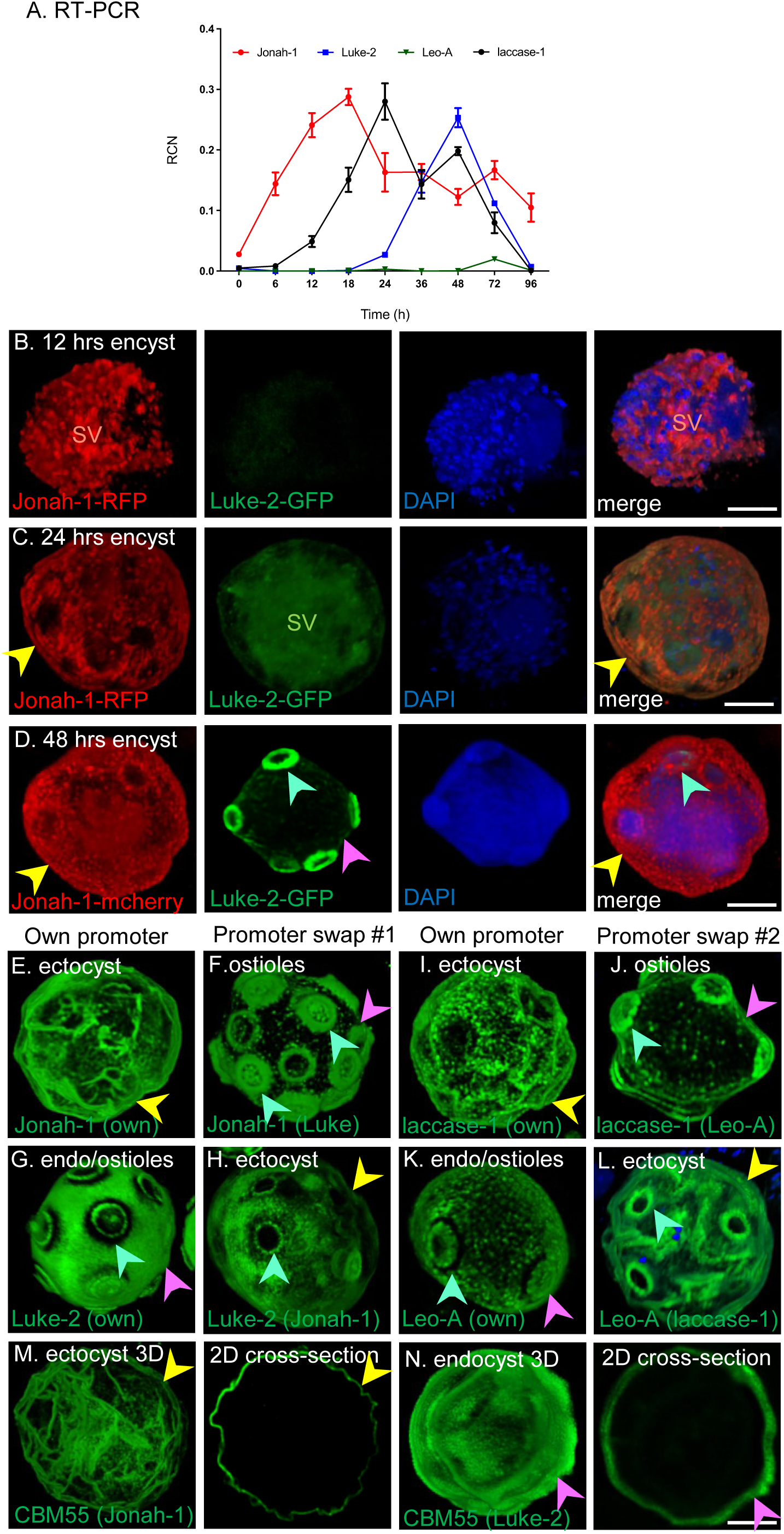
RT-PCR, double labels, promoter swaps, and chitin localization. (A) RT-PCR shows that ectocyst proteins (Jonah-1 and laccase-1) are made earlier than endocyst proteins (Luke-2 and Leo-A). Note values for each time points are normalized with calreticulin, which is highly expressed in trophozoites and encysting Ac. (B) After 12 hrs encystation, Ac are filled with secretory vesicles (SV) containing Jonah-1-RFP, as well as CFW-labeled vesicles most likely containing cellulose, while Luke-2-GFP is absent. (C) After 24 hrs encystation, Jonah-1-RFP localizes to the endocyst layer, while CFW and Luke-2-GFP are predominantly in secretory vesicles. (D) After 48 hrs encystation, the cyst wall has the appearance of mature cysts with Jonah-1-mCherry in ectocyst layer, Luke-2 GFP in ostioles and to a lesser extent the endocyst layer, and CFW predominantly in the endocyst layer. (E) In the first promoter swap, Jonah-1-GFP expressed under its own earlier promoter localizes to the ectocyst layer (yellow arrow) (see also Fig 1B). (F) Expression under the later Luke-2 promoter causes Jonah-1-GFP to localize to the ostioles (blue-green arrows), which increase in thickness and number, while there is minimal localization of Jonah-1-GFP to the endocyst layer (pink arrows). (G) Conversely, Luke-2-GFP under its own later promoter localizes to the endocyst layer and forms a narrow ring around ostioles (see also Fig 2B). (H) Expression under the earlier Jonah-1 promoter causes Luke-2 to relocate to the ectocyst layer. Remarkably, the narrow ring of Luke-2-GFP around the ostioles appears the same under either its own or the Jonah-1 promoter, suggesting its target glycopolymer is available early and later during encystation. (I) In the second promoter swap, laccase-1-GFP expressed under its own earlier promoter localizes to the ectocyst layer (see also Fig 4D). (J) Expression under the later Leo-A promoter causes laccase-1-GFP to localize to the ostioles with minimal localization to the endocyst layer. (K) Conversely, Leo-A-GFP under its own later promoter localizes to the endocyst layer and forms a narrow ring around ostioles. (L) Expression under the earlier laccase promoter causes Leo-A to localize to the ectocyst layer and forms a narrow ring around ostioles. These experiments show that under both early and late promoters, Jonah-1 and Laccase-1 have similar localizations in the cyst wall, suggesting that each bind to the same glycopolymer(s). Luke-2 and Leo-A also have similar localizations under early and late promoters, but the localization is distinct from that of Jonah-1 and Laccase-1, suggesting pairs of wall proteins are binding to different glycopolymers. (M) GFP-tagged CBM55 expressed under an earlier Jonah-1 promoter localizes to chitin fibrils in the ectocyst layer. (N) GFP-tagged CBM55 expressed under a later Luke-2 promoter localizes to a honey comb of chitin in the endocyst layer, which is distinct from the pattern of any GFP-tagged cell wall protein. Scale bars for B to N are each 5 µm.

Double labels on the same plasmid show Jonah-1-RFP is made at 12 hrs of encystation (Fig. 5B), when there is no evidence of Luke-2-GFP while CFW labels glycopolymers (most likely cellulose) that are present in small vesicles. After 24 hrs encystation (Fig. 5C), Jonah-1-RFP begins to accumulate on the single-layered wall, which is lightly labeled with Luke-2-GFP that is predominantly in small secretory vesicles that are not well resolved from each other. After 48 hrs encystation (Fig. 5D), Jonah-1-mCherry localizes to the ectocyst layer with a slightly punctate appearance, as shown in Fig 1B and 5E, while Luke-2-GFP forms rings around ostioles and lightly labels the endocyst layer, which is heavily labeled with CFW. This appearance of Jonah-1-mCherry, Luke-2-GFP, and CFW is indistinguishable from that of mature cysts at 72 or 96 hrs encystation (data not shown).

For the promoter swap experiments, we used GFP-tagged proteins that are either made earlier and localized to the ectocyst layer (Jonah-1 and laccase-1) or made later and localize to the endocyst layer and ostioles (Luke-2 and Leo-A) (Fig. 1B, 2B, 3D to 3F, and 4B to 4D) (29, 37, 38). In the first promoter swap, Jonah-1-GFP expressed under the later Luke-2 promoter moves from the ectocyst layer to the ostioles, which are densely coated and increase in number from ∼9 to as many as 16, while the endocyst layer is weakly labeled (Fig. 5E and 5F). The localization of Jonah-1-GFP under Luke-2 promoter is distinct from those of GFP-tagged Luke-2 and Leo-A under their own promoters, which show much greater localization to the endocyst layer, suggesting Jonah-1 binds to different glycopolymers (Fig. 1B and 2B). Conversely, Luke-2-GFP expressed under the earlier Jonah-1 promoter moves from the endocyst layer and ostioles to lightly decorate the ectocyst layer and form flat, narrow rings around the ostioles (Fig. 5G and 5H). These narrow rings, which match those of Luke-2-GFP under its own promoter, suggest that the glycopolymer in the ostiole to which Luke-2 is binding is available both earlier and later during encystation. The localization of Luke-2-GFP in the ectocyst layer is distinct from those of GFP-tagged Jonah-1, Jonah-3(c), Leo-S(c), or laccase-1, which, under their own promoters, coat the ectocyst layer and form dimples where ostioles are located (Fig. 1B, 1D, 3F, and 4D). This result suggests that glycopolymers bound by Luke-2 are likely not the same as those bound by proteins that normally localize to the ectocyst layer.

In the second promoter swap experiment, we expressed laccase-1-GFP (earlier) and Leo-A-GFP (later) each under its own promoter or the other promoter and determined their localizations in mature cyst walls. Laccase-1-GFP expressed under the later Leo-A promoter moves from the ectocyst layer to the ostioles, which are densely coated (Fig. 5I and 5J). Leo-A-GFP expressed under the earlier laccase-1 promoter, moves from the endocyst layer and ostioles to decorate lightly the ectocyst layer and form flat rings around the ostioles (Fig. 5K and 5L).

When expressed under the earlier Jonah-1 promoter, the chitin-binding domain of the *Entamoeba* cellulase (CBM55) tagged with GFP is localized to distinct fibrils in the ectocyst layer, some of which have linear arrays of small spheres (Fig. 5M). A 2D cross-section of the same cell shows that the chitin layer is narrow. In contrast, CBM55-GFP expressed on the later Luke-2 promoter has a honey-comb appearance in the endocyst layer, which is thicker on 2D cross-section, and there is no localization to ostioles (Fig. 2N).

In summary, qRT-PCR and double labels support our previous conclusions that Jonah-1 and laccase-1 in the ectocyst layer proteins are made earlier, while Luke-2 and Leo-A in the endocyst layer and ostioles are made later (29). Promoter swaps support our hypothesis that earlier expression does not just correlate with but causes proteins to localize to the ectocyst layer and later expression causes proteins to localize to the endocyst layer and ostioles. The similar localizations of Jonah-1 and laccase-1 under earlier and later promoters, which are distinct from localizations of Luke-2 and Leo-A under similar promoters, suggest pairs of wall proteins bind to distinct glycans. Finally, CBM55-GFP suggests chitin forms fibrils in the ectocyst layer and honeycombs in the endocyst layer, the mechanism of which is unknown but fascinating.

### Structures of abundant wall proteins reveal the complex ancestry of Ac cyst wall proteins and identify linear arrays of three aromatic amino acids in Luke-2 and Leo-A, which bind cellulose and direct proteins to the Ac cyst wall

Previously, we used sequence-based searches to show that Jonah lectins contain one or three choice of anchor-A (CAA) domains like a spore coat protein of *Bacillus anthracis*, while Luke lectins contain two or three domains similar to those of cellulose-binding proteins of *Dictyostelium discoideum* to carbohydrate-binding modules of bacteria (CBM2) and plant (CBM49) endocellulases (29, 32, 42–45, 54–57). Further, the Ac laccase-1 has three cupredoxin-like domains like those of bacterial and fungal enzymes, while 8-Cys domains of Leo appear to be unique to Ac (50, 51). Here we used AlphaFold and Foldseek to 1) predict structures of abundant cyst wall proteins and compare them to proteins with proven structures, 2) identify cellulose-binding sites for Luke and Leo lectins, and 3) select domains in Jonah-1 and laccase-1 for immunizing rabbits to produce antibodies that detect cysts by immunofluorescence microscopy (39–41).

Luke-2 and Luke-3 have two and three β-jelly-roll folds (BJRFs), respectively, each of which has a disulfide bond linking its beginning to its end. The BJRFs of Luke lectins are separated by short (33 to 47-aa) unstructured domains enriched in Ser (yellow) or Thr (blue) (Fig. 2D and 2E). Slimes molds including *Dictyostelium, Tieghemostelium, Heterostelium, Cavenderia*, and *Polysphondylium* also have cell wall proteins with one to five similar BJRFs, consistent their shared ancestry with Luke lectins (32, 56). Further, the BJRFs of Luke-2 and a predicted *Dictyostelium* cellulose-binding protein (Q86KB6_DICDI) each contain a linear array of three aromatic amino acids that are also present in CBM2 of an endocellulase of a cellulolytic soil bacterium *Cellulomonas fimi,* the structure of which has been solved (PDB 1EXH) (pink arrows in Fig. 2F and 2G) (58, 59). Ala mutations of these three aromatic amino acids in CBM2 and in CBM49 of a plant endocellulase, which is closely related, show they are the binding sites for cellulose (42, 47). Here we muted to Ala two sets of three aromatics (W35, W73, and W88 and W187, W228, and F244) of Luke-2, which was fused to maltose-binding protein (MBP) and expressed in the periplasm of *E. coli* where, like the endoplasmic reticulum of eukaryotic cells, disulfide bonds are formed (29, 60, 61). We purified MBP-Luke-2 +/- Ala mutations on amylose resins and tested their binding to Avicel (crystalline) cellulose. Western blots showed WT Luke-2 binds well to Avicel cellulose, while MBP alone and Luke-2 plus Ala-mutations fail to bind to cellulose (Fig. 2J). Further, Luke-2-GFP plus Ala mutations, which was expressed under the same promoter as WT Luke-2-GFP, no longer localizes in the endocyst layer and ostioles (Fig. 2B and 2C). The binding of MBP-WT Luke-2 to Avicel cellulose argues for its proper folding, so we used it to immunize rabbits.

Leo lectins have two sets of four disulfide knots (4DKs), which are adjacent (Leo-A) or separated by a long, unstructured, Thr-rich domain (Leo-S(c)) (Fig. 3G and 3I). Although there are numerous carbohydrate-binding modules composed of sets of 4DKs (e.g., CBM18 of WGA, CBM19 of a *Saccharomyces cerevisiae* GH18 chitinase, and CBM55 of *Entamoeba histolytica* GH18 chitinase), we were unable with Foldseek to identify any shared structures with the set of 4DKs of Leo (43, 46, 52, 62–64). Remarkably, 4DKs of Leo-A and Leo-S(c) contain linear arrays of three aromatic residues, even though these 4DKs have no relationship to BJRFs of Luke or to CBM2 and CBM49 of bacterial and plant endocellulases (Fig. 3H). Ala mutations to two sets of three aromatics (Y46, Y63, and Y77 and Y134, Y151, and Y165) caused an MBP-Leo-A fusion to no longer bind to Avicel cellulose (Fig. 2J). Similarly, the Ala mutant of Leo-A-GFP, which was expressed under the same promoter as WT Leo-A-GFP, no longer localizes to the endocyst layer and ostioles (Fig. 3B and 3C). As above, the binding of MBP-WT Leo-A to Avicel cellulose argues for its proper folding, so we used it to immunize rabbits. Jonah-1 has a single three-sided BHF, a pair of α-helices of unknown function, and a long, unstructured,

Thr-rich domain, while Jonah-3(c) has three BHFs and three long, unstructured, Ser-rich domains (Fig. 1E and 1F). The BHF of Jonah-1 closely resembles the three-sided BHF of an antifreeze protein of an Antarctic sea ice bacterium *Colwella sp*., the structure of which has been solved (PDB 3WP9) (Fig. 1E to 1H) (65). Although Jonah-1 and laccase-1 (next paragraph) contained numerous aromatic amino acids on their surface, none was in linear arrays, so we did not perform Ala mutations to test cellulose-binding. Finally, we chose the BHF of Jonah-1 to make an MBP-fusion for immunizing rabbits, because we wanted to avoid unstructured, Thr-rich regions, which might interfere with proper folding of the protein and/or contain *O*-linked glycans that would obscure antigenic sites.

The laccase-1 of Ac has three cupredoxin-like domains (CuRO-1, CuRO-2 and CuRO-3) but lacks unstructured, Ser- or Thr-rich domains, which distinguishes it from other ectocyst layer proteins (Jonah-1, Jonah-3(c), and Leo-S(c)) (Fig. 4E). Instead, laccase-1 has a positively charged loop between CuRO-2 and CuRO-3, which is also present in spore coat proteins of bacteria. Ac laccase-1 shares a 44% identity with spore coat protein A of *Caldicoprobacter faecalis*, which is present in sewage sludge. Ac laccase-1 shows slightly lower positional identities with laccases of archaea (e.g., 40% with *Halalkalicoccus paucihalophilus*, fungi (e.g., 38% identity with *Penicillium nalgiovense*), and plants (e.g., 36% with *Diphasiastrum complanatum* (36%). *Pelomyxa schiedti* (36% identity) is the only Amoebazoa with a laccase, which is absent from humans and most metazoa but are present in a few arthropods (e.g., 28% identity with *Rotaria sp.*). While the high positional identity with bacterial laccases suggests the Ac laccase-1 may be active, we did not determine what substrates are oxidized, and we did not knock it out to determine the phenotype caused. The CuRO-1 of laccase-1, which we chose to make an MBP-fusion for immunizing rabbits, closely resembles that of the spore coat protein A of *Bacillus subtilis*, the structure of which has been solved (PDB 4YVN) (Fig. 4F) (66). A possible concern here is the presence of a 10-aa sequence (NVYAGLAGFY) near the C-terminus, which is also present in laccases of some bacteria, archaea, fungi, and plants and so might lead to cross-reacting antibodies.

In summary, BJRFs of Luke lectins share recent common ancestry with wall proteins of slime molds and distant ancestry with CBM2 and CBM49 of bacterial and plant endocellulases, while 4DKs of Leo are unique. BHFs of Jonah lectins and three CuRO domains of laccase closely resemble those of bacterial proteins and so likely derive by horizontal gene transfer, which was not proven here. Although the structures of the BJRFs of Luke-2 and the set of 4DKs of Leo-A show no resemblance, the linear arrays of three aromatic amino acids, which are involved in binding cellulose and localizing proteins in the endocyst layer and ostioles, are the same, consistent with convergent evolution. Finally, while the BHF of Jonah-1, BJRFs of Luke-2, and 4DKs of Leo-A have no well-conserved sequences that might lead to cross-reacting antibodies with bacteria or fungi in eye scrapings, CuRO-1 of laccase-1 has a well-conserved 10-aa sequence that may be problematic.

### Rabbit antibodies to Jonah-1, Luke-2, Leo-A, and laccase-1 all efficiently detect cysts of 10 of 11 *Acanthamoeba* isolates, including five T4 genotypes that cause most cases of AK

The goals here were to 1) use rAbs to recombinant wall proteins to visualize native proteins by western blots and confocal microscopy and 2) test how well each antibody detected CFW-labeled cysts of 11 *Acanthamoeba* species/strains.

Rabbits were immunized with MBP-fused to WT Luke-2, WT-Leo-A, BHF of Jonah-1, and CuRO-1 of the laccase-1 in complete Freund’s adjuvant and boosted three times with incomplete Freund’s adjuvant at Cocalico Biologics, Inc., and rabbit IgGs were purified using Protein-A Sepharose (60, 61). Western blots and confocal microscopy showed that pre-bleeds from rabbits do not react with trophozoites or cysts of Ac (Fig. 6A). While rAbs to MBP-fusions with Jonah-1, Luke-2, Leo-A and laccase-1 do not bind to trophozoite proteins, each rAb binds well to cyst wall proteins of Neff strain of Ac. Anti-Jonah-1 rAbs bind to a protein of the expected size of ∼55-kDa, as well as to two lower mol wt bands. The latter may result from proteolytic cleavage of Jonah-1 before or during isolation of cyst proteins and/or by cross-reaction with smaller Jonah-1 proteins, which share multiple conserved domains. Anti-Luke-2 rAbs bind to a thick ∼50-kDa band, which is greater than the 27-kDa expected size of Luke-2. We suspect that the increased size of Luke-2 is caused by extensive glycosylation of its five *N*-glycan sites and ∼20 *O*-glycan sites in the low complexity, Ser-rich spacer (67). Anti-Leo-A rAbs bind to a thick ∼13-kDa band, which is slightly smaller than the expected size of 17-kDa, perhaps due to proteolytic cleavage. Finally, anti-laccase-1 rAbs bind to a 75-kDa band, which is slightly bigger than the expected size of 64-kDa, as well as to a less abundant 55-kDa band. Again, addition of *N*-glycans or *O*-glycans may explain the increase in size, while proteolytic cleavage may explain the lower mol wt band.

**FIG 6.**
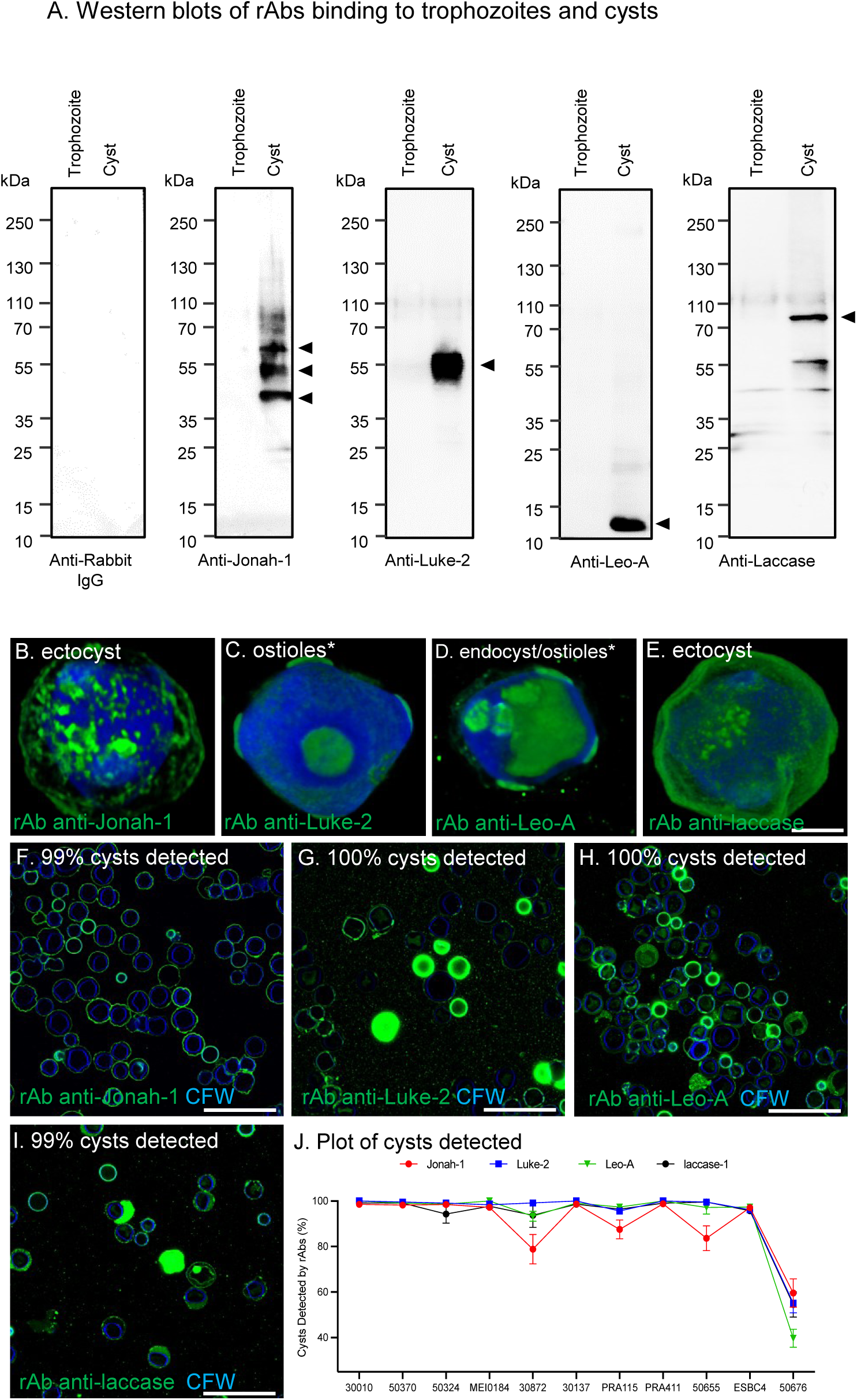
Binding of rAbs to cyst wall proteins and to Neff strain Ac (A) Western blots show pre-bleeds from rabbits (negative control) fail to bind to proteins of trophozoites or cysts, while rAbs fail to bind to trophozoite proteins (a second negative control) but bind well to cyst proteins of Neff strain. As discussed in detail in the Results, rAbs to the BHF of Jonah-1 binds to the expected 55-kDa band and to two lower mol wt bands. Anti-Luke-2 rAbs bind to a heavy and broad 50-kDa band, which is greater than the expected size of 27-kDa. Anti-Leo rAbs bind to a 13-kDa band, which is slightly smaller than the expected size of 17-kDA. Anti-laccase rAbs bind to a 75-kDa band, which is slightly bigger than the expected size of 64-kDa, as well as to a less abundant 55-kDa band. (B) High power confocal microscopy shows rAbs to Jonah-1 bind in a patchy distribution to the surface of Neff strain of Ac and other *Acanthamoeba* species, which are shown in Fig 7A. (C) Anti-Luke-2 rAbs bind strongly to ostioles and weakly to endocyst layer of some but not all Neff strain Ac. An asterisk is attached to “ostioles” because anti-Luke-2 rAbs can also bind to ectocyst layer, as shown in Fig 7B. (D) Anti-Leo-A rAbs also bind strongly to ostioles with a variable binding to the endocyst layer of some but not all of Neff strain Ac. Again, an asterisk indicates that anti-Leo-A rAbs can also bind to the ectocyst layer, as shown in Fig 7C. (E) Anti-laccase-1 rAbs bind in a homogenous manner to cysts of Neff strain, as well as to cyst of other Acanthamoeba species, which are shown in Fig 7D. (F to I). Low power confocal micrographs shows rAbs to Jonah-1, Luke-2, Leo-A and laccase-1 detect well CFW-labeled cysts of the Neff strain. (J) A plot shows the percentage of CFW-labeled cysts detected by four rAbs in two independent experiments (average plus SEM), in which 100+ cysts were counted in random low power fields. Rabbit antibodies to Luke-2, Leo-A, and Laccase-1 each detect >95% of cysts 10 of 11 *Acanthamoeba* isolates tested, while rAbs to Jonah-1 are slightly less efficient in detecting three isolates of *Acanthamoeba*. The exception is *A. mauritaniensis*, which was poorly detected with all four rAbs, suggesting a problem with cyst formation under the conditions used here. Single scale bar for B to E is 5 µm.

We tested rAbs versus the model Neff strain of Ac, as well as *Acanthamoeba* species/strains from the ATCC collected by Monica Crary, who studies their binding to contact lenses (24, 27). Segments of 18S RNA genes were amplified from trophozoites, cloned, and sequenced to confirm the genotype of 11 *Acanthamoeba* species/strains prior to encystation and testing with rAbs (13–16). Four T4 genotypes from the ATCC included *A. castellanii* Neff stain (30010), MA strain (50370), and unnamed strain (50324) and *A. polyphaga* (30872), while a fifth T4 (MEEI 0184) and another AK isolate (Esbc4) came from Noorjahan Panjwani, who discovered the mannose-binding protein on the surface of trophozoites (68). Other genotypes from ATCC included T7 (*A. astronyxis* 30137), T11 (*A. hatchetti* PRA-115) and T18 (*A. byersi* PRA-411 and *Acanthamoeba sp. 13* 50655).

High-power confocal microscopy showed rAbs to the BHF of Jonah-1 bound in a somewhat patchy distribution to the ectocyst layer (surface) of cysts of Neff strain and 10 other *Acanthamoeba* species and/or strains (Fig. 6A and 7A). This patchy distribution, which is in contrast to the homogeneous distribution of Jonah-1-GFP (Fig 1B), may result from masking of the BHF of Jonah-1 by its large, unstructured Thr-rich domain, which likely contains *O*-linked glycans, and/or masking by other ectocyst wall proteins (Fig. 1E). In contrast, rAbs to CuRO-1 of laccase-1 bound in a homogenous pattern to the Neff strain and to other *Acanthamoeba* species/strains, which is similar to that of laccase-1-GFP, suggesting there is less masking of laccase-1 (Fig. 4D, 6E, and 7D). While rAbs to Luke-2 and Leo-A densely labeled the ostioles and weakly labeled the endocyst layer of many cysts (Fig. 6C and 6D), other cysts were labeled on the endocyst layer and/or ectocyst layer (Fig. 7B and 7C). Of note, rAbs to Luke-2 and Leo-A often bound in similar patterns to a particular *Acanthamoeba* isolate, which is reminiscent of similar appearances of Luke-2-GFP and Leo-A-GFP in the promoter swap experiments (Fig. 5). There are two explanations, which are not mutually exclusive, for the variable distribution in the cyst walls of rAbs to Luke-2 and Leo-A. First, these proteins may be made at different times by different *Acanthamoeba* species/strains, so that Luke-2 and Leo-A localize to different places. Second, these rAbs may bind to relatively small amounts of Luke-2 and Leo-A in the ectocyst layer, which was not visualized by confocal microscopy, because of the large amounts of GFP-tagged Luke-2 and Leo-A in the endocyst layer and ostioles (Fig. 2B and 3B).

**FIG 7.**
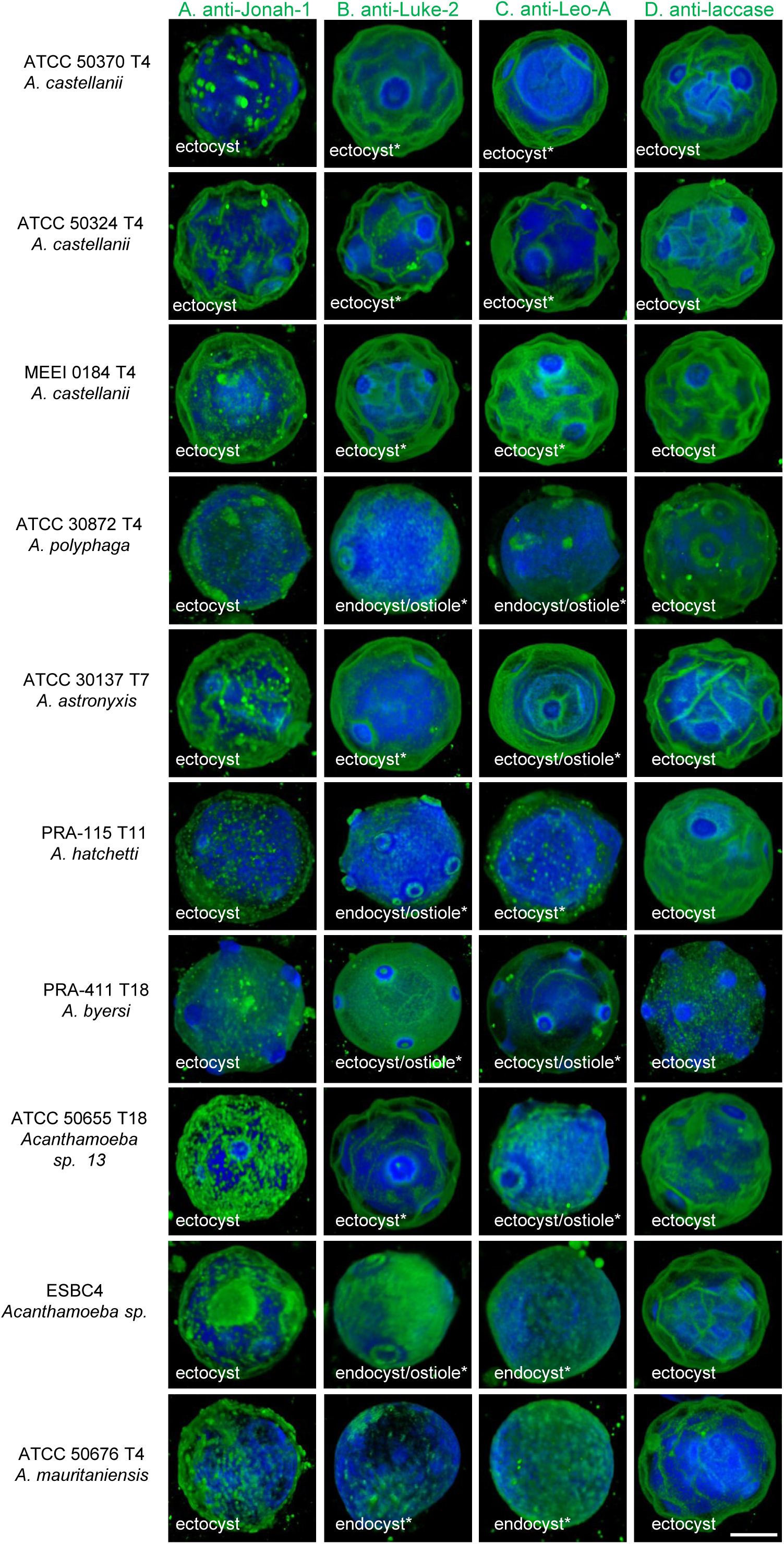
Binding of rAbs to Jonah-1, Luke-2, Leo-A, and Laccase to a diverse set of *Acanthamoeba Spp. C*ysts High power confocal micrographs show binding of rAbs to cysts ten *Acanthamoeba* isolates, the identities of which were confirmed by sequencing 18S RNAs. Column (A) Anti-Jonah-1 rAbs bind in a patchy distribution to the ectocyst layer (surface) of most cysts of *Acanthamoeba* isolates. Column (B) Anti-Luke-2 rAbs bind to the endocyst layer and ostioles of some cysts (e.g., *A. polyphaga, A. hatchetii, A, byersi*, and ESBC4) and to the ectocyst layer of other cysts (e.g., three *A. castellanii* and *A. astronyxis*). Asterisks indicate heterogeneous distribution of rAbs among the same *Acanthamoeba* isolate. Column (C) Anti-Leo-A rAbs also have a heterogeneous distribution on the ectocyst layer, endocyst layer, and/or ostioles. In some cases anti-Leo-A distribution matches that of anti-Luke-2 (e.g., *A. astronyxis, A. byersi*, and *A. sp. 13*), in the same way that Leo-GFP and Luke-GFP match in the promoter swap experiments (Fig 5). Column (D) Anti-Jonah-1 rAbs bind in a homogeneous distribution to the ectocyst layer (surface) of the vast majority of cysts of *Acanthamoeba* isolates. Counts of CFW-labeled cysts by each antibody are shown in Fig 6J. Single scale bar for A to D is 50 µm.

To determine which wall proteins might be the best targets for diagnosis of cysts in AK, we counted in randomly selected low power confocal fields the number of CFW-labeled cysts detected by rAbs to Jonah-1, Luke-2, Leo-A, and laccase-1 (Fig. 6F to 6I). In two separate experiments, >100 cysts were counted for each rAb and each *Acanthamoeba* species/strain, and the averages +/- SEM were plotted in Fig 6J. We found that rAbs to Luke-2, Leo-A, and laccase-1 each detected >95% of CFW-labeled cysts of 10 of 11 *Acanthamoeba* species and strains tested. In contrast, anti-Jonah-1 rAbs showed >95% detection for 7 of 11 *Acanthamoeba* species but detected somewhat fewer cysts of *A. byersi* (PRA-115) (T11) (91%), *Acanthamoeba* sp. 13 (ATCC 50655) (T18) (89%), and *A. polyphaga* (ATCC 30872) (T4) (85%). Why this is the case is not clear, but the result, which suggests that the BHF of Jonah-1 might not be quite as good a target as the BJRFs of Luke-2, the 4DKs of Leo-A, or CuRO-1 of laccase-1, was reproduced in separate experiments. In addition, all four rAbs struggled to detect *A. mauritaniensis* (T4) (ATCC 50676), suggesting that there is a problem with encysting these trophozoites under the conditions used here. While counts were performed for two sets of cysts labeled with Protein-A Sepharose-purified antibodies, we obtained similar results with rabbit sera diluted 1:300. Most important, we confirmed that rAbs to all four proteins were also visible with a conventional Zeiss fluorescence microscope, which is more similar to those present in clinical labs or in ophthalmologists’ offices. While the success with rAbs to Jonah-1 and laccase-1 was expected, because these proteins are on the ectocyst layer (surface) of cysts, the success of rAbs to Luke-2 and Leo-A was unexpected because we previously had trouble locating Luke-2-GFP or Leo-A-GFP using anti-GFP antibodies (29). Finally, regardless of whether the rAbs are binding to the ectocyst layer or endocyst layer, the presence of ∼10 ostioles makes *Acanthamoeba* cysts look distinct from fungal walls, which may at most have a few bud scars (29, 31, 34).

## DISCUSSION

### Incorrect protein prediction of Jonah-3 in AmoebaDB led to its artifactual but interesting localization in rings around ostioles

While Ac proteins sequences available in AmoebaDB made it possible for us to perform mass spectrometry of cyst walls and for others to discover dozens of proteins of interest, its predictions are based upon an incomplete genome of Neff strain of Ac and incomplete transcriptomes (17, 18, 29). We used an unfinished transcriptome to show TMHs between three BHFs, which incorrectly localized GFP-tagged Jonah-3-AmoebaDB to a ring around ostioles, were caused by frameshifts. Instead, Jonah-3(c) has low-complexity, Ser-rich spacers like those of Luke-2 and localizes to the ectocyst layer in a pattern matching that of Jonah-1. The unfinished transcriptome also allowed us to correct an eight amino acid long deletion in Leo-S-AmoebaDB that would have disabled its N-terminal 4DK, which is necessary for binding cellulose and localizing the protein to the ectocyst layer. Because AmoebaDB predicts 14,000 proteins, the errors described here in Jonah-3, Leo-S, and laccase-2 are mere anecdotes but suggest a need for a reannotated proteome based upon a well-constructed transcriptome. Finally, despite being an artifact, rings around the ostioles formed by Jonah-3-AmoebaDB may be useful for studying ostiole formation during encystation and ostiole breakdown during encystation. Similarly, the heavy labeling of ostioles by Jonah-1 and laccase-1 under the later Luke-2 and Leo-A promoters, respectively, may also facilitate studies of ostiole formation and degradation.

### Timing of expression and properties of abundant wall proteins determine their localization in the ectocyst layer, endocyst layer, and/or ostioles

Because whole cyst walls were purified by Percoll gradients and examined by mass spectrometry, we expressed selected proteins with a GFP-tag under their own promoters to precisely localize them in the two layered-wall connected by ostioles (29, 37, 38). Here we increased the number of ectocyst layer proteins from one (Jonah-1) to three (Jonah-3(c), Leo-S(c), and laccase-1). Laccase-1 and Leo-S(c) were also localized to a massive set of secretory vesicles early in encystation, as was previously shown for Jonah-1. For the first time, we used double labels to colocalize Jonah-1 and Luke-2 in developing walls and show unequivocally that the ectocyst layer is made first and the endocyst layer and ostioles are made second. Rabbit antibodies confirmed the presence of Jonah-1 and laccase-1 in the ectocyst layer of untransformed cysts of the model Neff strain of Ac, as well as cysts of 10 other *Acanthamoeba* species/strains. Promoter swaps showed Jonah-1 and laccase-1 switch to the endocyst layer and ostioles when expressed under later promoters of Luke-2 and Leo-A, respectively. Conversely, Luke-2 and Leo-A switch from the endocyst layer and ostioles to the ectocyst layer when expressed under the early promoters of Jonah-1 and laccase-1, respectively. Because the localizations of Jonah-1 and laccase-1 resemble each other under early and later promoters, it is likely that they are binding to the same glycopolymer(s), which we did not determine here. Similar localizations of Luke-2 and Leo-A under both promoters suggest these lectins are also binding to the same glycopolymer(s). This result is not surprising, as Luke-2 and Leo-A contain similar arrays of aromatic amino acids that bind cellulose. Finally, we took advantage of earlier and later promoters and a heterogenous probe from an *Entamoeba* chitinase to show two distinct appearances for chitin in the ectocyst layer (thick fibrils) and endocyst layer (honey comb), the latter of which we had never seen before with the exogenous probe for chitin (WGA) or GFP-tagged wall proteins.

Double labels are a promising tool for studying encystation by live protists, as well as studying how cyst walls are broken down by excysting Ac. Our conclusion that early expression causes proteins to localize in the ectocyst layer, and later expression causes proteins to localize in the endocyst layer and ostioles leaves many questions unanswered. As above, we do not know how ostioles are formed, and we do not know what causes chitin to have such distinct appearance in the two layers. In addition, we have not yet precisely localized cellulose in cyst walls or determined whether its appearance in the two layers is different. Finally, we have not identified sequences in regions 5’ to the start ATG that bind transcription factors specific for earlier or later encystation-specific expression, nor have we figured out why some encystation-specific promoters are stronger than others and so produce greater quantities of transcripts for wall proteins.

### While its 4DKs are unique, Leo lectins share many properties with wall proteins of Ac and other organisms, arguing for convergent evolution

Although the four families of Ac wall proteins show no relationship to each other, three have been previously identified and characterized in walls of protists, fungi, plants, and/or bacteria. Luke lectins have two or three BJRFs, which share recent ancestry with wall proteins of *Dictyostelium* and distant ancestry with CBM2 and CBM49 of bacterial and plant endocellulases (32, 42, 47, 56, 59). Laccases, which are most similar to those of bacteria, are widely distributed in walls of plants, fungi, and oomycetes (50, 51, 66). Although their function in Ac, which proliferate in temperate climates, is not clear, BHFs similar to those in Jonah lectins protect Artic bacteria from freezing (57, 65). While its 4DKs that bind cellulose are unique, Leo lectins share properties with wall proteins of Ac and other organisms, suggesting the importance of convergent evolution (69). Chitin-binding domains of *Saccharomyces* chitinases (CBM19), *Entamoeba* chitinases (CBM55), and WGA (CBM18) also contain unique 4DKs, while chitin-binding domains of Jacob lectins contain unique 3DKs (43, 46, 52, 63, 64). Even though 4DKs of Leo and BJRFs of Luke have no structural similarity, each contains linear arrays of three aromatic amino acids, which Ala mutations showed bind cellulose. Similar linear arrays of aromatics that bind cellulose are present in endocellulases of bacteria (CBM2) and plants (CBM49), as well as in CBM63s of expansins (proteins that unfold cellulose) of plants, bacteria, and Ac (42, 44, 47). Reinvention by convergent evolution of these linear arrays of aromatics strongly suggests they are the best means for binding cellulose, which forms flat ribbons with alternating glucose residues facing opposite surfaces.

Like Leo-A with two adjacent 4DKs and Leo-S with 4DKs separated by a large Thr-rich spacer, *Entamoeba* Jacob-1 has two adjacent 3DKs, while Jacob-2 has a third 3DK separated by a long, unstructured, Ser-rich spacer (70). Unstructured Ser- or Thr-rich domains, which are also present in Jonah and Luke lectins, are likely modified by *O*-linked glycans, which protect glycoproteins from being degraded by bacterial proteases, as shown for wall proteins of *Entamoeba* and *Cryptosporidium* (71, 72). Due to constraints on resources and time, we did not identify *O*-linked glycans on Ac cyst wall proteins, nor did we confirm the AlphaFold structures by crystallizing any of the four wall proteins studied here. This seems less of an issue for the BHFs of Jonah-1 and -3, BJRFs of Luke-2 and Luke-3, and CuRO-1 domain of laccase-1, the structures of which match those of crystallized proteins (59, 65, 66). A goal of future studies will be to solve the crystal structure of the unique 4DK of Leo +/- cellulose.

### Abundant wall proteins do not have to be in the ectocyst layer to be good targets for diagnostic antibodies

While it is rare for us (and other parasitologists) to pair basic and translational science experiments, it worked well here for the following reasons. The BHF of Jonah-1, the BJRFs of Luke-2, the 4DKs of Leo-A, and CuRO-1 domain of laccase-1 each expressed well as MBP-fusions in the periplasm of bacteria, where disulfide bonds are formed (60, 61). In contrast to anti-peptide rAbs to Jonah-1 and Leo-A, which bound to Western blots but not to fixed cysts (29), rAbs to MBP-fusions of all four Ac wall proteins bound well to Western blots and to cysts of Neff strain of Ac, as well as to cysts of 9 of 10 other isolates of *Acanthamoeba*. While we expected that rAbs to Jonah-1 and laccase-1, which are both in the accessible ectocyst layer, to efficiently detect CFW-labeled cysts, anti-laccase-1 performed slightly better than anti-Jonah-1. Even though we doubted whether rAbs to Luke-2 and Leo-A, which are in the relatively inaccessible endocyst layer and ostioles, would detect CFW-labeled cysts efficiently, anti-Luke-2 and anti-Leo-A rAbs both performed very well. Although the labeling of the ectocyst layer by rAbs to Luke-2 and Leo-A is not easy to explain, efficient detection is what matters for a diagnostic reagent. Finally, the signal from the secondary goat antibody binding to rAbs to Jonah-1, Luke-2, Leo-A, and laccase-1 is strong enough, so that cysts were easily detected with a conventional fluorescence microscope, not just by an expensive confocal microscope used to get three-dimensional, high-resolution images of the ectocyst layer, endocyst layer, and ostioles.

Because these rAbs cross-react with MBP that was not removed prior to vaccination, they are not ready for testing on corneal scrapings that likely contain numerous bacteria. Although we are confident that Luke-2, Leo-A, and Jonah-1 antigens are each unique to Ac, a 10-aa sequence in laccase-1 may lead to antibodies that cross-react with walls of bacteria, fungi, or plants. Cysts examined here were made by starving cultured *Acanthamoebae* and so might not be the same as those made in the corneal epithelium, soil, or water. Diagnostic anti-cyst antibodies will complement monoclonal antibodies to the mannose-binding proteins on trophozoites (68, 73), as well as antibodies to transporters and secreted proteins of trophozoites and cysts (74–76). Anti-cyst antibodies may also supplement Loop-mediated Isothermal Amplification (LAMP) assays for diagnosing AK (77, 78).

## MATERIALS AND METHODS

### Ethics statement

Culture and manipulation of *Acanthamoebae* under BSL-2 protocols were approved by the Boston University Institutional Biosafety Committee. Production of custom rabbit antibodies was approved by the Institutional Animal Care and Use Committee of Cocalico Biologics, Inc., Denver PA.

### Summary of new methods to study Ac wall proteins

Many methods are the same that we used in our mass spectrometric characterization of proteins in purified cyst walls of Ac, which were described in detail (29). New here is use of unfinished transcriptome to correct protein prediction in AmoebaDB for Jonah-3(c) and Leo-S(c). Confocal microscopy replaced structured illumination microscopy (SIM), because it is quicker and allowed us to examine many more parasites. RT-PCR and double labels with GFP and either RFP or mCherry were used to compare expression and localization of ectocyst and endocyst layer proteins, while an exogenous probe from an *Entamoeba* chitinase (CBM55) was used to localize chitin in ectocyst and endocyst layers. Two pairs of promoter swaps were used to show that localization in the two layers of the wall is not just correlated with timing of expression but is caused by timing of expression. AlphaFold was used to predict structures for each cyst wall protein and test with Ala mutations aromatic amino acids involved in binding cellulose. Foldseek was used to suggest the origin of domains in cyst wall proteins that could not be identified using sequence-based searches. Recombinant proteins, rather than peptides, were used to make rAbs to abundant wall proteins, which supported localizations of four GFP-tagged constructs under their own promoters. Binding of the rAbs to cysts of numerous *Acanthamoeba* genotypes, which were confirmed by PCR of 18S RNA genes, supported their use as targets for diagnostic antibodies.

### *Acanthamoeba* species and strains, culture, and cyst preparation

*A. castellanii* Neff strain (ATCC 30010) trophozoite parasites were obtained from the American Type Culture collection (ATCC). Trophozoites of other strains of *Acanthamoeba*, originally derived from human corneal infections and granulomatous encephalitis infections, were acquired from Dr. Monica Crary, Alcon Research, LLC, Fort Worth, TX, United States or from Noorjahan Panjwani of Tufts University Medical School (24, 68). All experiments were performed using *A. castellanii* Neff strain until otherwise mentioned.

Ac trophozoites were grown and maintained in axenic culture at 30°C in T-75 tissue culture flasks in 10 ml ATCC medium 712 (PYG plus additives) with antibiotics (Pen-Strep) (Sigma-Aldrich Corporation, St. Louis, MO) as described previously (21, 29, 79). Adherent log-phase trophozoites from stationary culture were detached with the help of a cell scraper and pelleted down by centrifugation at 500x g for 5 min followed by two washes with 1X phosphate buffered saline (PBS). Cysts were prepared from trophozoites by incubating them with encystation medium (EM, 20 mM Tris-HCl [pH 8.8], 100 mM KCl, 8 mM MgSO_4_, 0.4 mM CaCl_2_, and 1 mM NaHCO_3_) (29, 79). In brief, ∼10^7^ trophozoites obtained from a confluent flask were washed with 1x PBS and subsequently incubated with EM in a T-75 tissue culture flask for 120 hours. The mature cysts were released with the help of a cell scrapper, harvested, and washed with 1x PBS by centrifugation at 1,500x g for 10 min. The harvested cysts were either used immediately or stored at 4°C or -20°C depending upon the need of the experiment.

### *In silico* sequence analysis of candidate cyst wall lectins

The full-length coding sequence of different cyst wall proteins of *A*. *castellanii* namely Jonah-1 (ACA1_16481), Jonah-3 (ACA1_157320), Luke-2 (ACA1_377670), laccase-1 (ACA1_068450), Leo-A (ACA1_074730) and Leo-S (ACA1_188350) were obtained from AmoebaDB, a functional genomic database useful for genetic studies of the Neff strain and ten other *Acanthamoeba* strains (https://amoebadb.org/amoeba/app) (80). The AmoebaDB database was also used to predict introns and identify paralog proteins and upstream promoters (nucleotide sequence) of different cyst wall proteins. The full-length amino acid sequences of candidate wall proteins were further analyzed, reannotated, and corrected based on homologs proteins using BLAST server in other species (54). Other *in-silico* tools used to functionally characterize the cyst wall proteins include the CDD database (conserved domain identification) (81), SignalP 4.1 (49) and DeepTMHMM (82) (signal peptides and transmembrane helices), CAZy and InterPro databases (Carbohydrate-binding modules) (43, 46, 55) (48) and big-PI (Glycosylphosphatidylinositol anchors) (83), respectively. Finally, we used an unfinished transcriptome of trophozoites and encysting Ac, which will be described elsewhere when completed, to check and correct protein predictions.

### *In silico* structural prediction of candidate cyst wall proteins

The 3D structure models of different cyst wall lectins were predicted by AlphaFold2 using the Colab server (https://colab.research.google.com/github/sokrypton/ColabFold/blob/main/AlphaFold2.ipynb) (39). The molecular visualization of active site residues, 3D structures, and structural superimposition was performed with the help of PyMOL Molecular Graphics System, Version 2.0 Schrödinger, LLC (84). Additionally, the predicted 3D structures were used to perform structural comparison and similarity search with large structure sets (AlphaFold database, PDB database, etc.) using Foldseek server (https://search.foldseek.com/search) (40).

### Expression and localization of candidate cysts wall lectins during encystation in *A. castellanii*

RT-PCR studies were performed to check the expression of cyst wall lectins in encysting protists. In brief, adherent log-phase trophozoites of Ac were washed twice with PBS and subsequently stimulated to encyst by incubation at 30°C with encystation media. The amoeba cells were collected at various time points including 0, 6, 12, 18, 24, 36, 48, 72, and 96 hours post incubation. The cells were pelleted down by centrifugation, washed with PBS, and stored in Trizol reagent at -80°C for further use. Once all the time points were collected, the Trizol cell pellets were thawed on ice, and the cells were lysed by bead beating (BioSpec mini bead beater) operated in a cold room at 4°C with 1 min beating and cooling off for 5 mins for 3 cycles. Total RNA was extracted using Directzol RNA minprep kit (Zymo Research), and the concentration was checked using Nanodrop 2000/2000c (Thermo Fisher Scientific). cDNA was synthesized using AMV reverse transcriptase (New England Biolabs, Ipswich, MA) as per the manufacturer instructions. The expression of abundant cyst wall proteins during encystation was analyzed using Realtime-PCR (Biorad) using Power SYBR green PCR master mix (Applied Biosystems). Calreticulin was used as housekeeping gene to normalize the expression profile of various candidate cyst wall lectins. The primers used for real time PCR analysis are listed in Supporting Information S1.

To verify the expression and subcellular localization of cyst wall proteins during encystation, we expressed individual cyst wall lectins from an episomal plasmid in Neff strain of Ac under their own promoter. Primers for making constructs are listed in Supporting Information S1, while sequences of promoters and proteins are listed in Supporting Information S2. For all constructs, the pGAPDH plasmid was used, which harbors a neomycin resistance gene (for G418 drug selection) and a glyceraldehyde 3-phosphate dehydrogenase (GAPDH) promoter for constitutive expression of a C-terminus GFP fusion chimera (37). We also used the pGAPDH vector to express proteins tagged with either RFP or mCherry, both codon-optimized for Ac. In our *in silico* structural studies, we observed a comparable binding site topography for Luke-2 and Leo-A cellulose-binding module (CBM), which consists of a linear array of aromatic amino acids, as previously observed phenomenon for other CBMs. To confirm the involvement of these aromatics in Luke-2 and Leo-A CBMs these aromatic amino acid residues were mutated to alanine. Tryptophan (W35, W73, W88, W187, W228 and phenylalanine (F244) of Luke-2 were replaced with alanine, while tyrosine (Y46, Y63, Y66, Y134, Y151, and Y165) of Leo-A were replaced with alanine. The construct used for Jonah-1, Leo-A (WT), and Luke-2 (WT) were taken from a previously published study (29). While the full-length coding sequences of Leo-A mutant, Luke-2 mutant, laccase-1, and Leo-S gene were codon optimized and custom synthesized from the Twist Biosciences along with their respective promoter sequences. For the promoter swap experiment, the Luke-2 promoter was replaced with Jonah-1, while the laccase-1 promoter was replaced with the Leo-A promoter and vice versa. For the chitin-binding domain CBM55 construct, *Entamoeba histolytica* CBM55 sequence was obtained from AmoebaDB, codon optimized for Ac, and expressed under Jonah-1 or Luke-2 promoter. For visualizing two proteins in the same cell, a single pGAPDH vector was engineered to express Jonah-1 tagged with RFP or mCherry and Luke-1-GFP, each expressed under its own promoter. RFP and mCherry were codon-optimized for Ac and synthesized at Twist Biosciences. The 5’ upstream sequence (400-600 bp) (Supporting Information S2) for each gene was PCR amplified from Ac genomic DNA (used as a promoter to drive their expression) to swap the respective gene promoter in the pGAPDH plasmid. For construct preparation, we used a restriction-free cloning strategy using NEBuilder HiFi DNA assembly master Mix from New England Biolabs, Ipswich, MA. The identities of final constructs were sequence verified using Oxford nanopore sequencing technology from Plasmidsaurus. The primers used for cloning of different promoters, and cyst wall lectin genes are listed in Supporting Information S1.

### Transfection

Transfection in *A. castellanii* was performed routinely in the lab using Lipofectamine™ Transfection Reagent (Thermo Fisher Scientific), as per the manufacturer’s instruction. Briefly, for each transfection ∼5 x 10^5^ log-phase trophozoites were seeded in a T-25 flask in ATCC medium 712 and incubated for 30 min at 30°C. After the incubation, the adherent trophozoites were washed, and the media was replaced with 500 μl of encystation media (EM). Prior to transfection, the Lipofectamine^TM^ 3000 reagent was diluted in EM (7.5 µL Lipofectamine^TM^ 3000 was diluted in 125 µL EM). The master mix of DNA was prepared by diluting 8 μg of plasmid DNA with EM (to achieve a final volume of 117 µL) and subsequent addition of 8 µL P3000^TM^ reagent. Diluted Lipofectamine^TM^ 3000 and DNA master mix both were added in a 1:1 (125 + 125 µl) ratio and incubated at room temperature for an additional 15 min. Following incubation, the DNA-lipofectamine complex (250 µL) mixture was added directly onto the adherent trophozoites in 1 mL EM (1250 µL). The trophozoites were further incubated for an additional 4 hours at 30°C. After this, the EM and transfection mixture was removed by pipetting and 10 ml fresh media was added into each well. After 24h of transfection, the used medium was replaced with fresh media containing G418 (12.5 µg/mL). After 48h of transfection, cells were transferred from 6 well plates to T-75 flasks with ATCC medium 712 plus G418 (25 µg/mL). The used medium was changed routinely with fresh media containing G418 (25 µg/mL) on every fourth day. After 2 to 4 weeks, the transfectants began growing robustly in the presence of G418.

### Confocal microscopy

To check the expression of the abundant cyst wall proteins during the encystation, the transgenic Ac trophozoites (which express GFP-tagged chimera cyst wall proteins) were induced to encyst with EM. The encysting amoebas were collected at various time points (0, 6, 12, 18, 24, 36, 48, 72, and 96 hours), centrifuged, washed (1x PBS), and fixed by incubating them with 4% paraformaldehyde (PFA) for 15 min at room temperature. Following fixation cells were pelleted down, washed, and resuspended in PBS. The fixed cells were stained with Wheat Germ Agglutinin 1:10 (WGA, 1 mg/ml, ThermoFisher Scientific) conjugated to Alexa Flour 647 and Calcofluor White 1:20 (CFW, 1mg/ml, Sigma Aldrich) in PBS for 30 min at room temperature. The cells were washed 3 times with 1x PBS and mounted in VECTASHIELD® Antifade Mounting Medium (Vector Laboratories, Newark, CA). Samples were illuminated using 380 nm (CFW), 488 nm (GFP chimera), and 647 nm (WGA) laser excitation. Alternatively, 647 nm laser excitation was used for RFP or mCherry chimeras. The fluorescence images were captured using CFI Plan Apochromat VC 60XC WI Plan APO 60x oil objective of Nikon Ni2 AX inverted confocal microscope equipped with NIR imaging system. We deconvolved 0.1 μm optical sections using NIS elements (Version: AR5.41.02) imaging software. All confocal images shown were 3D reconstructions using dozens of z-stacks. Size bars were based upon 2D cross-sections.

### Overexpression and purification of candidate cyst wall proteins as MBP-fusions

The MBP-fusion constructs of Jonah-1, laccase-1, Luke-2 (WT and mutant), and Leo-A (WT and mutant) were prepared by cloning the codon-optimized (*E.coli* expression) synthetic gene fragments into pMAL-p2x vector (New England Biolabs) for periplasmic expression in BL21-CodonPlus(DE3)-RIPL (Agilent Technologies, Lexington, MA) (61). The primers used for cloning synthetic gene fragments and the length of different domains of cyst wall proteins (devoid of signal sequence) are listed in Supporting Information S1 and S2. The expression of each MBP-fusion proteins was induced by the addition of 0.1 mM of IPTG for 16 h at 16°C. The bacterial cell pellet was lysed, and supernatant was collected as per the manufacturer’s specifications (New England Biolabs). AKTA start was used to purify the MBP tagged protein using an MBPtrap HP column (Cytiva). The MBPtrap column was equilibrated with equilibration or wash buffer (20 mM Tris-HCl (pH 8), 200 mM NaCl, 1 mM EDTA). Later the supernatant mixture was passed through the column for the MBP-tagged protein to bind to the column at 0.5 mL/min flow rate. The column washed with 20 column volumes of wash buffer, and the purified protein was eluted with elution buffer (20 mM Tris-HCl (pH 8), 200 mM NaCl, 1 mM EDTA, 10 mM maltose). The identity and purity of recombinant purified MBP-fusion proteins were confirmed by SDS-PAGE and Western immunoblotting analyses. The wild-type recombinant Jonah-1, Luke-2, Leo-A, and laccase-1 MBP-fusions were used to raise custom polyclonal antibodies (Cocalico Biologicals). Total IgG was purified from plasma samples of pre-immune and post-immunized rabbits via Protein A affinity chromatography (Pierce™ Protein A Agarose, Thermo Fisher Scientific, USA) as per manufacturer instructions. In brief, for Protein A Sepharose purification, plasma was diluted 2-fold with binding buffer (1x Tris-Buffered Saline, pH 7.4) and loaded onto a column containing Protein A Agarose beads. The diluted plasma was passed through the column twice, and beads were washed with 1x TBS (20-fold column volume). The bound IgG was eluted with 0.1 M glycine-HCl (pH 2.7), into the neutralizing buffer (1 M Tris-HCl, pH 9.0) and concentrated, and the buffer exchanged into PBS using a 30 kDa Amicon Ultra centrifugal filter (Millipore, USA).

### Cellulose binding assay using WT and Mutant Luke-2 and Leo-A

To check the cellulose binding activity of WT and mutant Luke-2 and Leo-A, we performed an *in vitro* cellulose binding assay. In this assay, recombinant purified MBP fusion Luke-2 (WT and Ala mutant) and Leo-A (WT and Ala mutant) proteins were used. A total of 1 μg MBP-fusion protein (in 100 μl of 1% NP40) was incubated with 0.5 μg Avicel microcrystalline cellulose (Sigma-Aldrich) for 3 hours at 4°C with rocking. Following binding, microcrystalline cellulose fibers were pelleted down by centrifugation, while the supernatant was collected in a separate tube (unbound fraction). The microcrystalline cellulose fibers (bound fractions) were washed three times with 1% NP40. The input material (total) unbound (U), and bound (B) fractions were boiled in the SDS sample buffer. Soluble proteins were separated on SDS-PAGE (4-15%), blotted to nitrocellulose membranes, blocked in 5% BSA, and detected using anti-MBP antibodies (New England Biolabs). As a negative control, microcrystalline beads were incubated with MBP alone.

### Detection of candidate cyst wall lectins in *A. castellanii* trophozoites and cysts

To detect the cyst wall proteins in Ac, log-phase trophozoites parasites and 120 h post-encysting mature cysts were harvested and lysed in SDS sample buffer. The lysates were separated on SDS-PAGE gel (4-15%), transferred on nitrocellulose membrane, and blocked in 5% BSA in PBS. The blots were probed with primary rabbit polyclonal antibodies (1:5000)/purified rabbit IgG (1:1000) raised against the different abundant cyst wall proteins. Anti-rabbit HRP conjugated IgG (Thermo Fisher Scientific) was used as the secondary antibody. Rabbit pre-immune serum or anti-rabbit IgG were used as control. Super Signal West Pico PLUS (Thermo Fisher Scientific) substrate was used for chemiluminescent detection. Blots were imaged using GE ImageQuant LAS 4000 gel imager.

### Immuno-staining of the mature cysts with purified rabbit polyclonal antibodies

To check the applicability of custom raised polyclonal rabbit antibodies against detection of Jonah-1, Luke-2, Leo-A, and laccase paralogs, we performed IFA imaging using mature cysts from other *Acanthamoeba* species/strains. For IFA imaging, ∼0.5 to 1.0 x 10^7^ mature cysts (120 h post encystation) from different strains of *Acanthamoeba* were washed in PBS and fixed in 4% paraformaldehyde for 15 minutes at room temperature. Following fixation, cysts were washed three times with PBS and blocked with 1% BSA for 1 h at room temperature. The cysts were further incubated with the rabbit polyclonal antibody (1:200 dilution) for 1h at room temperature. Subsequently these cysts were washed three times with PBS and incubated with secondary anti-rabbit IgG conjugated with Alexa flour 488 (1:300) and Calcofluor White 1:20 (CFW, 1 mg/ml, Sigma Aldrich) for 30 min at room temperature. The cysts were washed 3 times with 1x PBS and mounted in VECTASHIELD® Antifade Mounting Medium (Vector Laboratories, Newark, CA). Samples were illuminated using 380nm (CFW), and 488 nm (Alexa flour 488) laser excitation. The stained cysts were imaged using confocal microscope as detailed above. For each rAb and each *Acanthamoeba* isolate, we counted at least 100 cysts in random fields to determine what percentage of CFW-labeled cysts were detected with the rAb. This experiment was repeated twice and counts for the two experiments were averaged. Finally, we examined the same set of slides using a Plan-APO Chromat 100x oil immersion lens on a Zeiss Axio Observer Z1 microscope equipped with Axio Cam ERc5s to show that a confocal microscope was not needed to detect rAb-labeled cysts.

## ACKNOWLEDGMENTS

Thanks to Jonathan Stefely (Massachusetts General Hospital), Sarah Calvo (Broad Institute and Howard Hughes Medical Institute), and Brian Haas (Broad Institute) for access to the unfinished transcriptome analysis of *Acanthamoeba castellanii* Neff strain. Thanks to Monica Crary of Alcon, Inc. and Noorjahan Panjwani of Tufts University for cultures of *Acanthamoeba*. Thanks to Cataldo Leone (Dean of BUSDM) for financial support for this work. This paper is dedicated to Jude, the cousin of Jonah, Luke, and Leo.

Supporting Information S1 lists the primers used for making constructs and for performing qRT-PCR. Supporting Information S2 lists promoter sequences and coding sequences of the abundant cyst wall proteins used for making GFP-tagged proteins. Also listed are the coding sequences used to make MBP-fusions for testing cellulose binding and for immunizing rabbits.

